# Development of a highly effective African swine fever virus vaccine by deletion of the I177L gene results in sterile immunity against the current epidemic Eurasia strain

**DOI:** 10.1101/861666

**Authors:** Manuel V. Borca, Elizabeth Ramirez Medina, Ediane Silva, Elizabeth Vuono, Ayushi Rai, Sarah Pruitt, Lauren G. Holinka, Lauro Velazquez Salinas, James Zhu, Douglas P. Gladue

## Abstract

African swine fever virus (ASFV) is the etiological agent of a contagious and often lethal disease of domestic pigs that has significant economic consequences for the swine industry. The disease is devastating the swine industry in Central Europe and East Asia, with current outbreaks caused by circulating strains of ASFV derived from the 2007 Georgia isolate (ASFV-G), a genotype II ASFV. In the absence of any available vaccines, African Swine Fever (ASF) outbreak containment relies on control and culling of infected animals. Limited cross protection studies suggest that in order to ensure a vaccine is effective it must be derived from the current outbreak strain or at the very least from an isolate with the same genotype. Here we report the discovery that deletion of a previously uncharacterized gene, I177L, from the highly virulent ASFV-G produces complete virus attenuation in swine. Animals inoculated intramuscularly with the virus lacking the I177L gene, ASFV-G-ΔI177L, in a dose range of 10^2^ to 10^6^ HAD_50_ remained clinically normal during the 28 day observational period. All ASFV-G-ΔI177L-infected animals had low viremia titers, showed no virus shedding, developed a strong virus-specific antibody response and, importantly, they were protected when challenged with the virulent parental strain ASFV-G. ASFV-G-ΔI177L is one of the few experimental vaccine candidate virus strains reported to be able to induce protection against the ASFV Georgia isolate, and the first vaccine capable of inducing sterile immunity against the current ASFV strain responsible for recent outbreaks.

**Importance:** Currently there is no commercially available vaccine against African swine fever. Outbreaks of this disease are devastating the swine industry from Central Europe to East Asia, and they are being caused by circulating strains of African swine fever virus derived from the Georgia2007 isolate. Here we report the discovery of a previously uncharacterized virus gene, which when deleted completely attenuates the Georgia isolate. Importantly, animals infected with this genetically modified virus were protected from developing ASF after challenge with the virulent parental virus. Interestingly, ASFV-G-ΔI177L confers protection even at low doses (10^2^ HAD_50_) and remains completely attenuated when inoculated at high doses (10^6^ HAD_50_), demonstrating its potential as a safe vaccine candidate. At medium doses (10^4^ HAD_50_) sterile immunity is achieved. Therefore, ASFV-G-ΔI177L is a novel efficacious experimental ASF vaccine protecting pigs from the epidemiologically relevant ASFV Georgia isolate.

## Introduction

African Swine Fever (ASF) is a contagious and often fatal viral disease of swine. The causative agent, ASF virus (ASFV), is a large enveloped virus containing a double-stranded (ds) DNA genome of approximately 190 kilobase pairs (1). ASFV shares aspects of genome structure and replication strategy with other large dsDNA viruses, including the *Poxviridae, Iridoviridae* and *Phycodnaviridae* (2). However, on a protein or amino acid level there is little homology with the majority of the viral proteins, and very few ASFV proteins have been evaluated for their functionality or for their contribution to virus pathogenesis.

Currently, ASF is endemic in more than twenty sub-Saharan African countries. In Europe, ASF is endemic on the island of Sardinia (Italy) and outbreaks in additional countries began with an outbreak in the Caucasus region in 2007, affecting Georgia, Armenia, Azerbaijan and Russia. ASF has continued to spread uncontrollably across Europe and Asia with ASFV outbreaks occurring in 2018-2019 in China, Mongolia, Vietnam, Laos, Cambodia, Serbia, Myanmar, North Korea and the Philippines. ASF has also spread to wild boar in Belgium, but has been restricted to a quarantine zone since the first introduction of the disease in 2018. Sequencing of several contemporary epidemic ASFVs suggests high nucleotide similarity with only minor modifications compared to the initial 2007 outbreak strain, ASFV Georgia 2007/1, a highly virulent isolate that belongs to the ASFV genotype II group (3).

There is no vaccine available for ASF and disease outbreaks are currently quelled by animal quarantine and slaughter. Attempts to vaccinate animals using infected cell extracts, supernatants of infected pig peripheral blood leukocytes, purified and inactivated virions, infected glutaraldehyde-fixed macrophages, or detergent-treated infected alveolar macrophages failed to induce protective immunity (4–7). Protective immunity develops in pigs surviving viral infection with moderately virulent or attenuated variants of ASFV, with long-term resistance to homologous, but rarely to heterologous, virus challenge (8, 9). Significantly, pigs immunized with live attenuated ASF viruses containing genetically engineered deletions of specific ASFV virulence-associated genes are protected when challenged with homologous parental virus (10–15). These observations constitute the only experimental evidence describing the rational development of an effective live attenuated vaccine against ASFV.

Here we report the discovery that deletion of a previously uncharacterized gene, I177L, from the highly virulent ASFV Georgia isolate (ASFV-G) results in complete attenuation in swine. Animals inoculated with the virus lacking the I177L gene, ASFV-G-ΔI177L, remained clinically normal, developed a strong virus-specific antibody response and, importantly, ASFV-G-ΔI177L-infected swine were completely protected when challenged with the virulent parental ASFV-G.

## Materials and Methods

### Cell culture and viruses

Primary swine macrophage cell cultures were prepared from defibrinated swine blood as previously described (15). Briefly, heparin-treated swine blood was incubated at 37°C for 1 hour to allow sedimentation of the erythrocyte fraction. Mononuclear leukocytes were separated by flotation over a Ficoll-Paque (Pharmacia, Piscataway, N.J.) density gradient (specific gravity, 1.079). The monocyte/macrophage cell fraction was cultured in plastic Primaria (Falcon; Becton Dickinson Labware, Franklin Lakes, N.J.) tissue culture flasks containing macrophage media, composed of RPMI 1640 Medium (Life Technologies, Grand Island, NY) with 30% L929 supernatant and 20% fetal bovine serum (HI-FBS, Thermo Scientific, Waltham, MA) for 48 hours at 37°C under 5% CO_2_. Adherent cells were detached from the plastic by using 10 mM EDTA in phosphate buffered saline (PBS) and were then reseeded into Primaria T25, 6- or 96-well dishes at a density of 5×10^6^ cells per ml for use in assays 24 hours later.

Comparative growth curves between ASFV-G and ASFV-G-ΔI177L viruses were performed in primary swine macrophage cell cultures. Preformed monolayers were prepared in 24-well plates and infected at a MOI of 0.01 (based on HAD_50_ previously determined in primary swine macrophage cell cultures). After 1 hour of adsorption at 37°C under 5% CO_2_ the inoculum was removed and the cells were rinsed two times with PBS. The monolayers were then rinsed with macrophage media and incubated for 2, 24, 48, 72 and 96 hours at 37°C under 5% CO_2_. At appropriate times post-infection, the cells were frozen at <-70°C and the thawed lysates were used to determine titers by HAD_50_/ml in primary swine macrophage cell cultures. All samples were run simultaneously to avoid inter-assay variability.

Virus titration was performed on primary swine macrophage cell cultures in 96-well plates. Virus dilutions and cultures were performed using macrophage medium. Presence of virus was assessed by hemadsorption (HA) and virus titers were calculated by the Reed and Muench method (16).

ASFV Georgia (ASFV-G) was a field isolate kindly provided by Dr. Nino Vepkhvadze, from the Laboratory of the Ministry of Agriculture (LMA) in Tbilisi, Republic of Georgia.

### Microarray analysis

The microarray data of ASFV open reading frames were obtained from the data set deposited in NCBI databases by Zhu et al. (submitted). In brief, total RNA was extracted from primary swine macrophage cell cultures infected with ASFV Georgia strain or mock infected at 3, 6, 9, 12, 15, and 18 hours post-infection (hpi). A custom designed porcine microarray manufactured by Agilent Technologies (Chicopee, MA) was used for this study. Both infected and mock-infected RNA samples were labeled with Cy3 and Cy5 using an Agilent low-input RNA labeling kit (Agilent Technologies). Cy5-labeled infected or mock-infected samples were co-hybridized with Cy3-labeled mock-infected or infected samples in one array, respectively, for each time point using a dye-swap design. The entire procedure of microarray analysis was conducted according to protocols, reagents and equipment provided or recommended by Agilent Technologies. Array slides were scanned using a GenePix 4000B scanner (Molecular Devises, San Jose, CA) with the GenePix Pro 6.0 software at 5 μm resolution. Background signal correction and data normalization of the microarray signals and statistical analysis were performed using the LIMMA package. The signal intensities of ASFV open reading frame RNA were averaged from both Cy3 and Cy5 channels.

### Construction of the recombinant ASFV-G-ΔI177L

Recombinant ASFVs were generated by homologous recombination between the parental ASFV-G genome and recombination transfer vector by infection and transfection procedures using swine macrophage cell cultures (15). Recombinant transfer vector (p72mCherryΔI177L) containing flanking genomic regions to amino acids 112 though 150 of the I177L gene, mapping approximately 1kbp to the left and right of these amino acids, and a reporter gene cassette containing the mCherry gene with the ASFV p72 late gene promoter, p72mCherry, was used. This construction created a 112 bp deletion in the I177L ORF (Fig. 1). Recombinant transfer vector p72mCheryΔI177L was obtained by DNA synthesis (Epoch Life Sciences Missouri City, TX, USA).

**Fig. 1.**
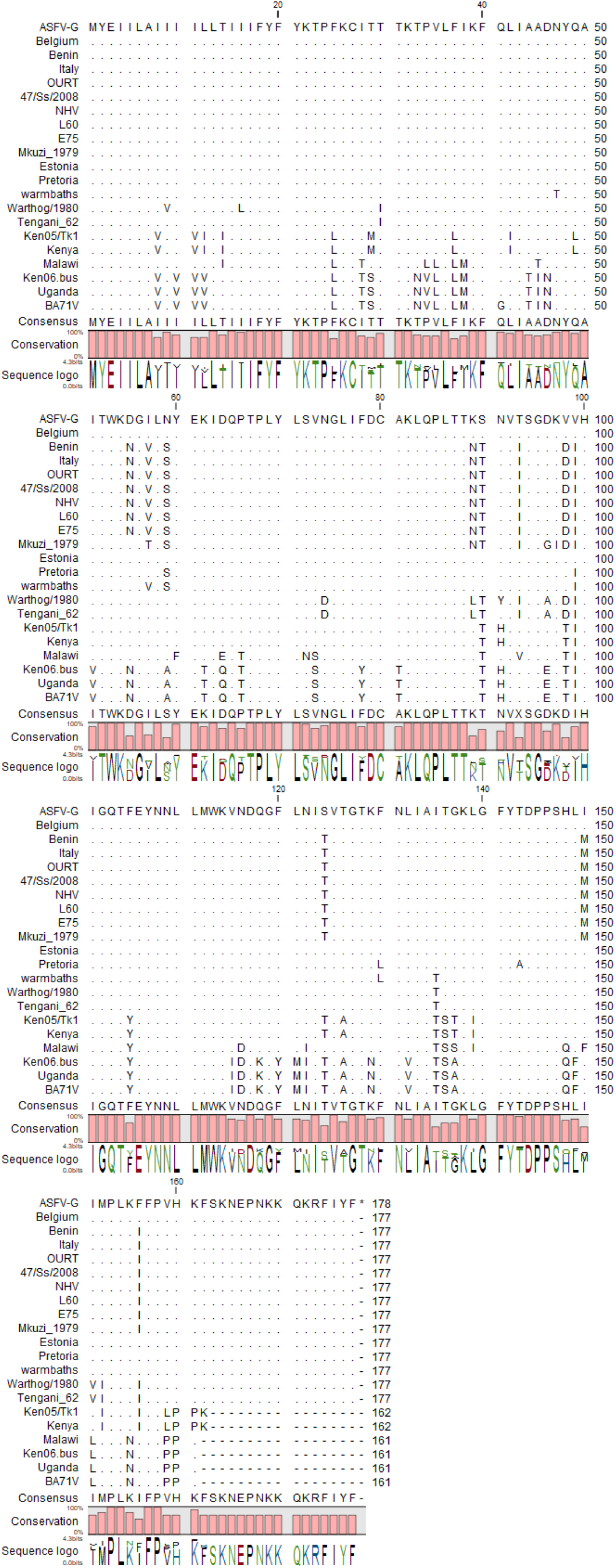
Multiple sequence alignment of the indicated ASFV isolates of viral protein I177L, matching residues are represented as ‘.’ and gaps in the sequence are represented by Degree of conservation between the sequences is represented below the sequences.

### Next Generation Sequencing (NGS) of ASFV genomes

ASFV DNA was extracted from infected cells and quantified as described earlier. Full-length sequence of the virus genome was performed as described previously (17) using an Illumina NextSeq500 sequencer.

### Animal experiments

Animal experiments were performed under biosafety level 3AG conditions in the animal facilities at Plum Island Animal Disease Center (PIADC) following a protocol approved by the PIADC Institutional Animal Care and Use Committee of the US Departments of Agriculture and Homeland Security (protocol number 225.04-16-R, 09-07-16).

ASFV-G-ΔI177Lwas assessed for its virulence phenotype relative to the virulent parental ASFV-G virus using 80-90 pound commercial breed swine. Groups of pigs (n=5) were inoculated intramuscularly (IM) either with 10^2^ - 10^6^ HAD_50_ of ASFV-G ΔI177L or 10^2^ HAD_50_ of parental ASFV-G virus. Clinical signs (anorexia, depression, fever, purple skin discoloration, staggering gait, diarrhea and cough) and changes in body temperature were recorded daily throughout the experiment. In protection experiments, animals inoculated with ASFV-GΔI177L were 28 days later IM challenged with 10^2^ HAD_50_ of parental virulent ASFV-G strain. Presence of clinical signs associated with the disease was recorded as described earlier.

### Detection of anti-ASFV antibodies

ASFV antibody detection used an in-house indirect ELISA, developed as described previously (18). Briefly, ELISA antigen was prepared from ASFV-infected Vero cells. Maxisorb ELISA plates (Nunc, St Louis, MO, USA) were coated with 1 μg per well of infected or uninfected cell extract. The plates were blocked with phosphate-buffered saline containing 10% skim milk (Merck, Kenilworth, NJ, USA) and 5% normal goat serum (Sigma, Saint Louis, MO). Each swine serum was tested at multiple dilutions against both infected and uninfected cell antigen. ASFV-specific antibodies in the swine sera were detected by an anti-swine IgM or IgG-horseradish peroxidase conjugate (KPL, Gaithersburg, MD, USA) and SureBlue Reserve peroxidase substrate (KPL). Plates were read at OD630 nm in an ELx808 plate reader (BioTek, Shoreline, WA, USA). Sera titers were expressed as the log10 of the highest dilution where the OD630 reading of the tested sera at least duplicates the reading of the mock infected sera.

## Results

### Conservation of I177L gene across different ASFV isolates

ASFV-G ORF I177L encodes for a 177 amino acid protein and is positioned on the reverse strand between nucleotide positions 174471 and 175004 of the ASFV-G genome (Fig. 1). The degree of I177L conservation among ASFV isolates was examined by multiple alignment using CLC genomics workbench software (CLCBio; Aarhus, Denmark). The ASFV I177L protein sequences were derived from all sequenced isolates of ASFV representing African, European and Caribbean isolates from domestic pig, wild pig, and tick sources. I177L has a predicted protein length of 161 to 177 amino acid residues depending on the isolate. Most of the isolates contain a protein with a length of 177 amino acids, with a few isolates showing a truncated C-terminus that yield a protein of 161-162 amino acids. It is predicted that I177L contains a possible N-terminal transmembrane helix (data not shown). I177L is sometimes annotated in recent isolates using a different start codon that occurs at position 112, however this has been recently shown to be a sequencing mistake in the annotation of these genomes(19). I177L at the amino acid level revealed a very high degree of amino acid identity among isolates when compared to isolates containing the same or different forms of I177L (Fig. 1).

### I177L gene is transcribed as a late gene during the virus replication cycle

The time of transcription of the I177L gene was determined by microarray evaluating total RNA extracted from primary swine macrophage cell cultures infected with ASFV-G at 3, 6, 9, 12, 15, and 18 hours hpi (representing an approximate one life cycle of ASFV replication).

Figure 2 shows the microarray signal intensities of three ASFV open reading frames at the six time points sampled. CP204L gene (encoding for ASFV protein p30) was expressed at approximate 41k photons per pixel at 3 hpi, which agrees with its known early gene expression after ASFV infection. The expression gradually decreased at 6 and 9 hpi and then significantly increased by more than 9-fold, reaching a plateau at 12 hpi. In contrast, B646L, a p72 virus capsid protein gene known for its late expression, was practically not expressed at 3 hpi at a level less than 20 photons per pixel. The p72 expression significantly increased to >44k at 6 hpi and reached a plateau at 12 hpi. I177L appears to be a late expressed gene much like B646L. The I177L gene was transcribed at less than 50 photons per pixel at 3 hpi. Unlike the p72 gene, I177L expression increased linearly at a much slower rate and its expression remained low at 18 hpi.

**Fig. 2.**
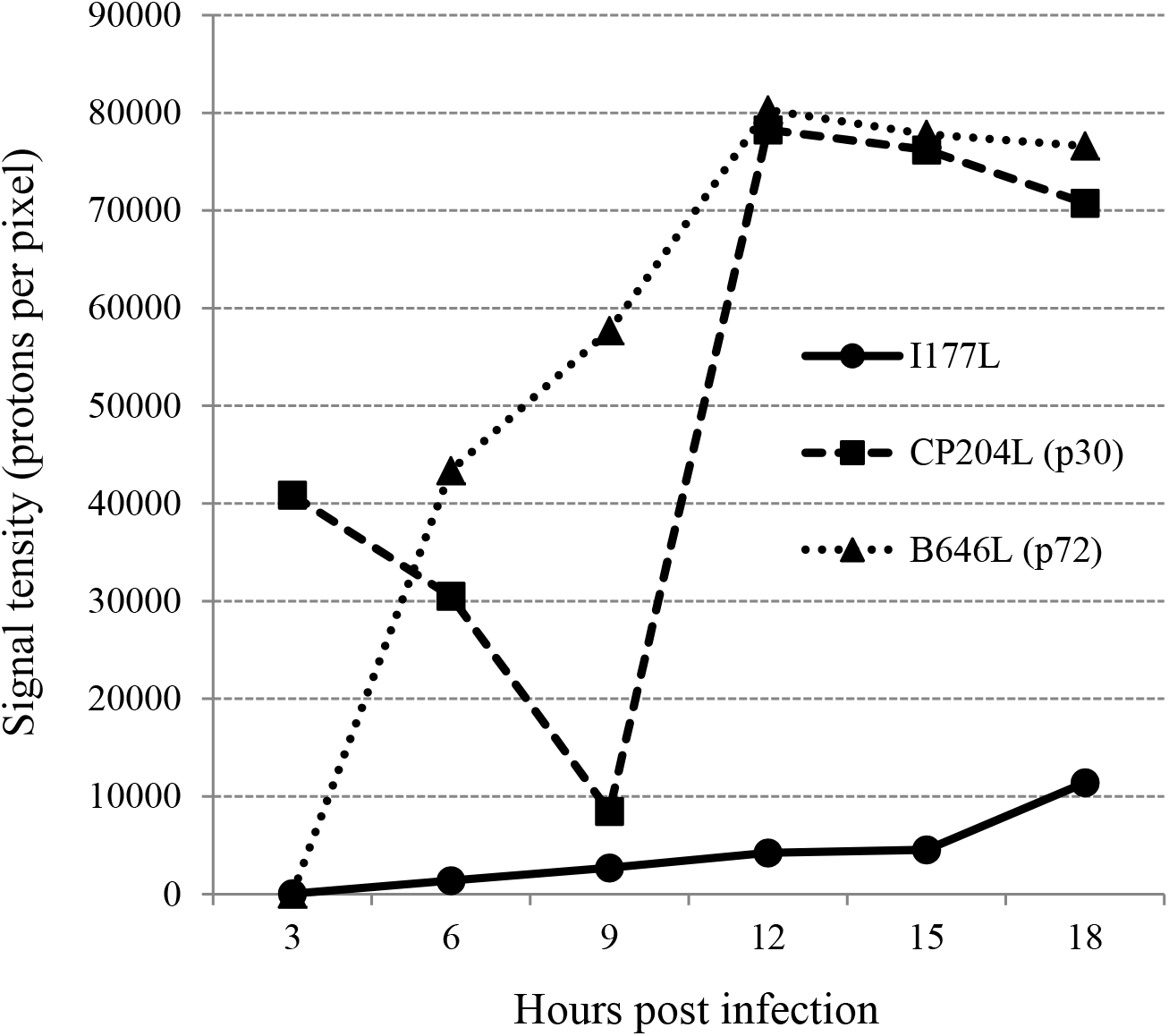
Time course of I177L gene transcriptional activity. Averaged microarray signal intensities (photons per pixel) of ASFV I177L, CP204L and B646L open reading frame RNA prepared from *ex vivo* pig macrophages infected with ASFV at 3, 6, 9, 12, 15 and 18 hours post infection.

### Development of the I177L gene deletion mutant of the ASFV-Georgia isolate

To determine the role of I177L during ASFV infection in cell cultures and virulence in swine, a recombinant virus lacking the I177L gene was designed (ASFV-G-ΔI177L). ASFV-G-ΔI177L was constructed from the highly pathogenic ASFV Georgia 2010 (ASFV-G) isolate by homologous recombination procedures as described in Material and Methods. The I177L gene was replaced by a cassette containing the fluorescent gene mCherry under the ASFV p72 promoter (Fig. 3). Recombinant virus was selected after 10 rounds of limiting dilution purification based on the fluorescent activity. The virus population obtained from the last round of purification was amplified in primary swine macrophage cell cultures to obtain a virus stock.

**Fig. 3.**
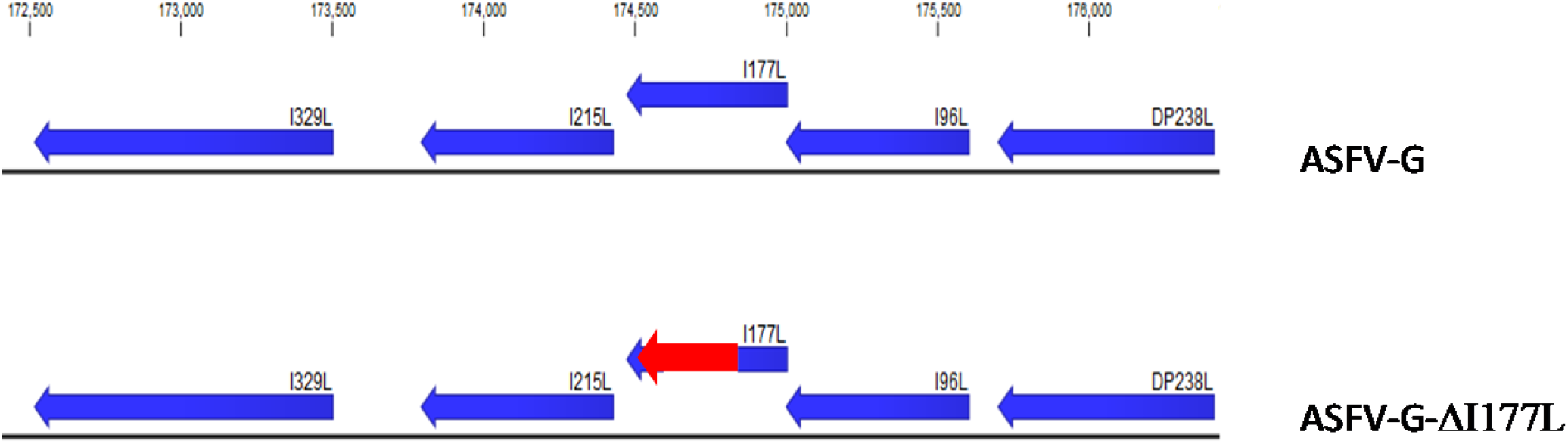
Diagram indicating the position of the I177L open reading frame in the ASFV-G genome. The donor plasmid with the homologous arms to ASFV-G and the mCherry under control of the p72 promoter in the orientation as indicated (in red). The final genomic changes introduced to develop ASFV-Georgia-ΔI177L where the sequence of the donor plasmid mCherry reporter is introduced to replace the ORF of I177L as indicated.

To evaluate the accuracy of the genetic modification, the integrity of the genome and the purity of the recombinant virus stock, full genome sequences of ASFV-G-ΔI177L and parental ASFV-G were obtained using Next Generation Sequencing (NGS) for comparison. The DNA sequence assemblies of ASFV-G-ΔI177L and ASFV-G revealed a deletion of 112 nucleotides (between nucleotide positions 174,530-174,671) from the I177L gene corresponding with the introduced modification. The consensus sequence of the ASFV-G-ΔI177L genome showed an insertion of 3,944 nucleotides corresponding to the p72mCherryΔI177L cassette sequence introduced within the 112 nucleotide deletion in the I177L gene. Besides the designed changes, no unwanted additional mutations were detected in the rest of the ASFV-G-ΔI177L genome.

### Replication of ASFV-G-ΔI177L in primary swine macrophages

*In vitro* growth characteristics of ASFV-G-ΔI177L were evaluated in primary swine macrophage cell cultures, the primary cell targeted by ASFV during infection in swine and compared relative to parental ASFV-G in multistep growth curves (Fig. 4). Cell cultures were infected at a MOI of 0.01 and samples were collected at 2, 24, 48, 72 and 96 hours post-infection (hpi). Results demonstrated that ASFV-G-ΔI177L displayed a growth kinetic significantly decreased when compared to parental ASFV-G. ASFV-G-ΔI177L yields are approximately 100 to 1,000-fold lower than those of ASFV-G depending on the time point considered. Therefore, deletion of the I177L gene significantly decreased the ability of ASFV-G-ΔI177L, relative to the parental ASFV-G isolate, to replicate *in vitro* in primary swine macrophage cell cultures.

**Fig. 4:**
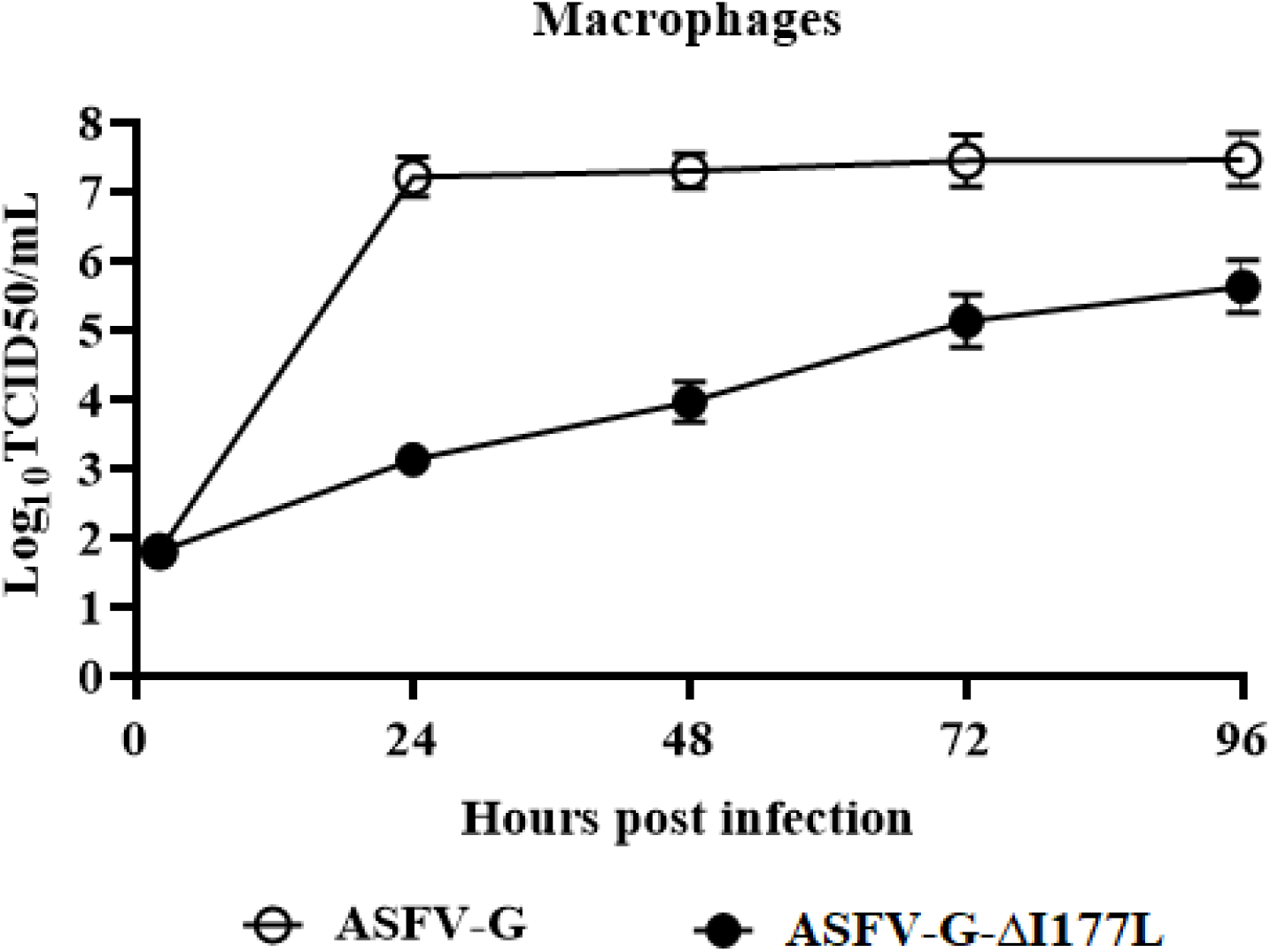
*In vitro* growth characteristics of ASFV-Georgia-ΔI177L and parental ASFV-G. Primary swine macrophage cell cultures were infected (MOI=0.01) with each of the viruses and virus yield titrated at the indicated times post-infection. Data represent means from three independent experiments. Sensitivity of virus detection: > 1.8 log_10_ HAD_50_/ml.

### Assessment of ASFV-G-ΔI177L virulence in swine

To evaluate the degree of attenuation reached by ASFV-G-ΔI177L, a preliminary experiment was performed using low virus load. A group of 80-90 pounds pigs were inoculated via IM with 10^2^ HAD_50_ ASFV-G-ΔI177L and compared with animals inoculated with 10^2^ HAD of parental ASFV-G. All five animals inoculated with ASFV-G had increased body temperature (>104 ^°^F) by day 5 postinfection, presenting with clinical signs associated with the disease including anorexia, depression, purple skin discoloration, staggering gait and diarrhea (Table 1). Signs of the disease aggravated progressively over time and animals were euthanized *in extremis* by 7 days post-infection (pi). Conversely, the five animals inoculated via IM with ASFV-G-ΔI177L did not present with any ASF-related signs, remaining clinically normal during the entire 28-day observation period.

**Table 1:** Swine survival and fever response following infection with 10^2^ HAD_50_ doses of ASFV-G-ΔI177L or parental ASFV-G.

| Virus and dose (HAD<sub>50</sub>) | No. of survivors/<br>total | Mean time to death<br>(days ± SD) | Fever | | |
| --- | --- | --- | --- | --- | --- |
| | | | No. of days to onset<br>(days ± SD) | Duration No. of days<br>(days ± SD) | Maximum daily temp<br>(°F ± SD) |
| ASFV-G | 0/5 | 7 (0)<sup>(1)</sup> | 5 (0) | 2 (0) | 106.1 (1.38) |
| ASFV-G-ΔI177L | 5/5 | - | - | - | 102.6 (0.56) |

(1) All animals were euthanized due to humanitarian reasons following the corresponding IACUC protocol.

Animals infected with ASFV-G presented with expected high homogenous titers (10^7.5^ - 10^8.5^ HAD_50_/ml) on day 4 pi, increasing (around 10^8.5^ HAD_50_/ml) by day 7 pi when all animals were euthanized. Conversely, ASFV-G-ΔI177L revealed a different pattern with low viremia (10^1.8^ – 10^5^ HAD_50_/ml) at day 4 pi, reaching peak values (10^4^ - 10^7.5^ HAD_50_/ml) by day 11 pi and then decreasing titers (10^2.3^ - 10^4^ HAD_50_/ml) until day 28 pi (Fig. 5). It should be noted that one of the five animals inoculated with ASFV-G-ΔI177L showed a remarkably lower viremia (1,000- to 10,000-fold lower depending on the time point considered) than the average viremia values of the other animals in the group. Therefore, deletion of the I177L gene produced complete attenuation of the parental highly virulent ASFV-G virus when inoculated at a low dose, with the infected animals presenting long viremias with relatively low values.

**Fig. 5:**
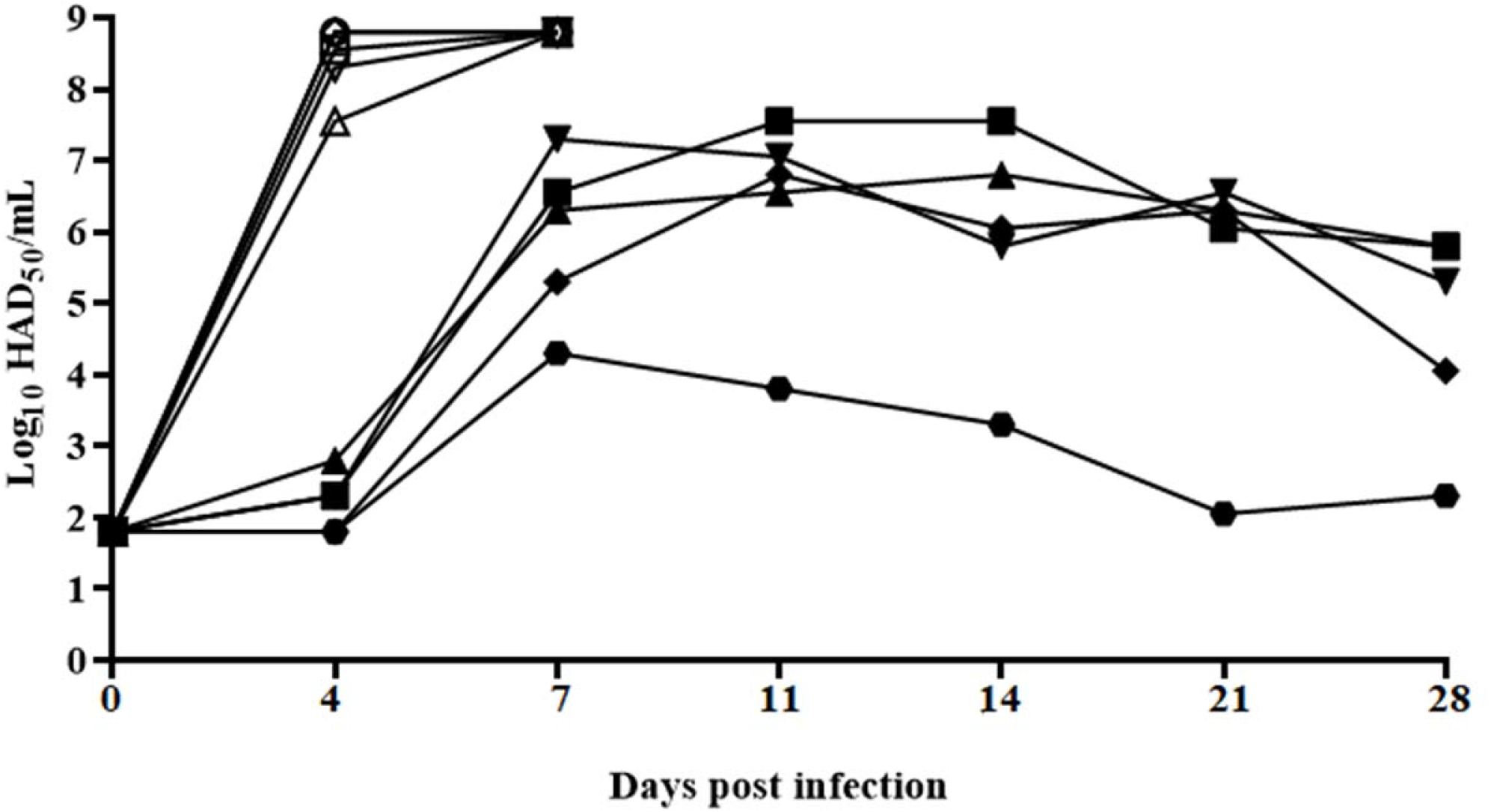
Viremia titers detected in pigs IM inoculated with either 10^2^ HAD_50_ of ASFV-Georgia-ΔI177L (filled symbols) or 10^2^ HAD_50_ of ASFV-G (empty symbols). Each curve represents values from individual animals in each group. Sensitivity of virus detection: ≥ 1.8 log_10_ HAD_50_/ml.

To investigate the potential presence of residual virulence in ASFV-G-ΔI177L a second experiment was performed where different groups of five pig were infected IM with either 10^2^, 10^4^, or 10^6^ HAD_50_ of ASFV-G-ΔI177L and their behavior compared with that of naïve animals inoculated with 10^2^ HAD_50_ of parental ASFV-G. In addition, a mock infected animal cohabitated in each of the groups, acting as a sentinel to detect potential virus shedding from the infected animals.

As in the first experiment, animals inoculated with ASFV-G exhibited all typical clinical signs of the disease and were euthanized *in extremis* by day 6-7 pi (Table 2). Interestingly all animals inoculated with ASFV-G-ΔI177L, including those receiving 10^6^ HAD_50_, did not present any ASF-related signs, remaining clinically normal during the entire observation period (28 days). Similarly, all sentinel animals remained clinically normal (Fig. 6).

**Fig. 6:**
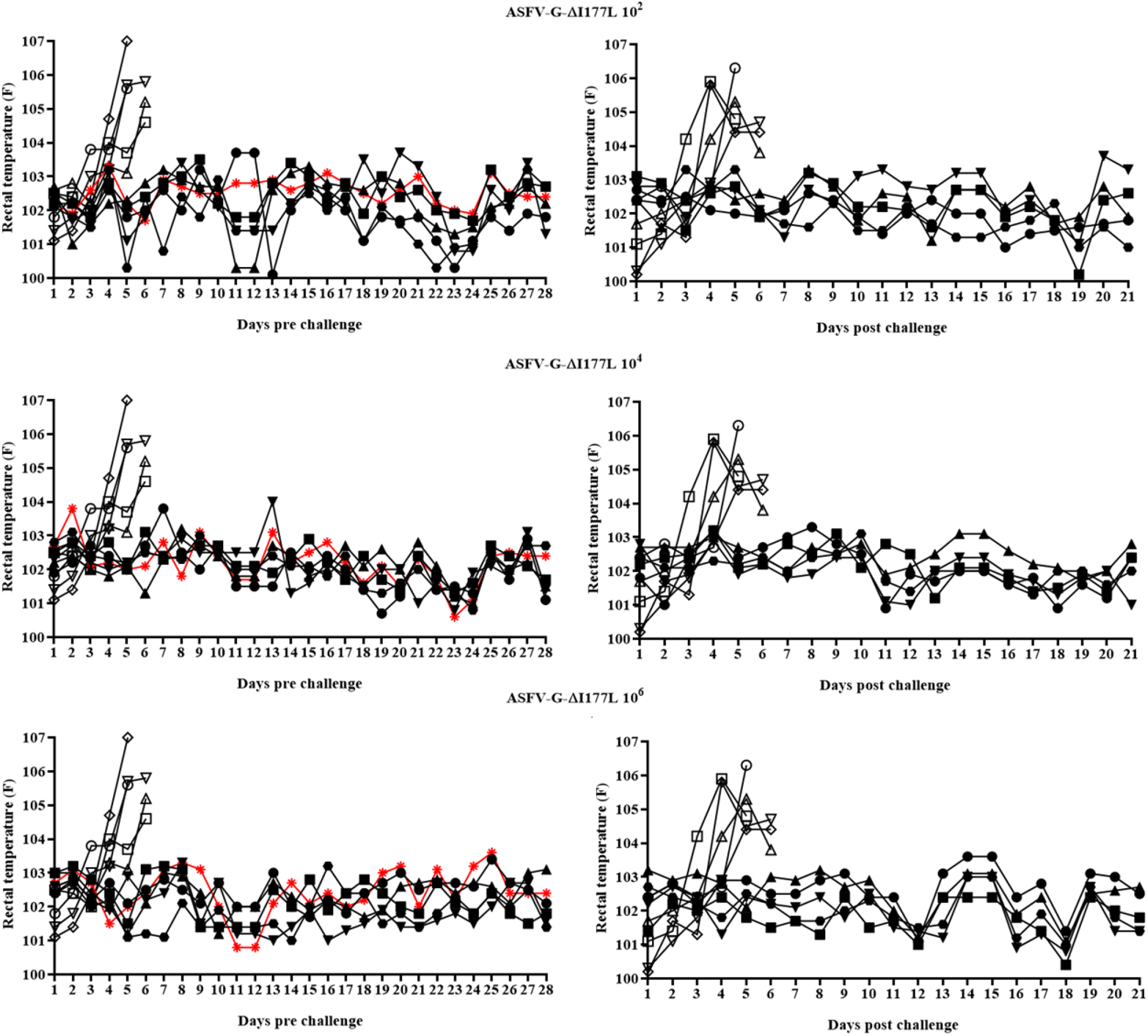
Kinetics of body temperature values in pigs IM inoculated with either 10^2^, 10^4^, or 10^6^ HAD_50_ of ASFV-Georgia-ΔI177L (filled symbols), mock inoculated (sentinels, showed in red) or 10^2^ HAD_50_ of ASFV-G (empty symbols) **(panels on the left)**, and after the challenge with 10^2^ HAD_50_ of ASFV-G **(panels on the right).** Each curve represents individual animals values in each of the group.

**Table 2.** Swine survival and fever response following infection with different doses of ASFV-G-ΔI177L or parental ASFV-G.

| Virus | No. of survivors/<br>total | Mean time to death<br>(days ± SD) | Fever | | |
| --- | --- | --- | --- | --- | --- |
| | | | No. of days to onset<br>(days ± SD) | Duration No. of days<br>(days ± SD) | Maximum daily temp<br>(°F ± SD) |
| ASFV-G | 0/5 | 6.6 (0.55)<sup>(1)</sup> | 4 (0) | 2.6 (0.55) | 105.2 (0.6) |
| ASFV-G-ΔI177L $10^2$ HAD<sub>50</sub> | 5/5 | - | - | - | 102.9 (0.5) |
| ASFV-G-ΔI177L $10^4$ HAD<sub>50</sub> | 5/5 | - | - | - | 102.8 (0.57) |
| ASFV-G-ΔI177L $10^6$ HAD<sub>50</sub> | 5/5 | - | - | - | 102.8 (0.49) |

(1) All animals were euthanized due to humanitarian reasons following the corresponding IACUC protocol.

Viremia kinetics in ASFV-G-infected animals presented high titers (10^3.5^ - 10^8^ HAD_50_/ml) on day 4 pi increasing (around 10^7.5^ HAD_50_/ml) by day 6-7 pi when all animals were euthanized (Fig. 7). Animals infected with 10^2^ HAD_50_/ml of ASFV-G-ΔI177L showed similar results to those seen in the previous experiment although this time viremias were not detected until day 11 pi (with the exception of one animal) and two out of the five animals presented significantly lower titers than the other three animals in the group. In the groups of animals infected with 10^4^ or 10^6^ HAD_50_/ml of ASFV-G-ΔI177L, viremias were clearly detectable at 4 days pi with average values remarkably higher (1,000-to 10,000-fold) than the group inoculated with 10^2^ HAD_50_/ml particularly, at 4 and 7 days pi. Heterogeneity in the viremia measurements is also seen in the groups inoculated with the higher doses of ASFV-G-ΔI177L particularly at 21 and 28 days pi when 3 animals in each group presented remarkably lower titers than the other two animals in the group. Therefore, deletion of the I177L gene produced a complete attenuation of the parental highly virulent ASFV-G virus even when used at high dosage, with the infected animals presenting low-titer viremias that persisted throughout the duration of the 28-day observational period. Interestingly, no virus was detected in any of the samples (all sampled blood time points as well as tonsil and spleen samples obtained at 28 days pi) obtained from sentinel animals (data not shown), indicating that ASFV-G-ΔI177L-infected animals did not shed enough virus to infect naïve pigs during the 28 days of cohabitation.

**Fig. 7:**
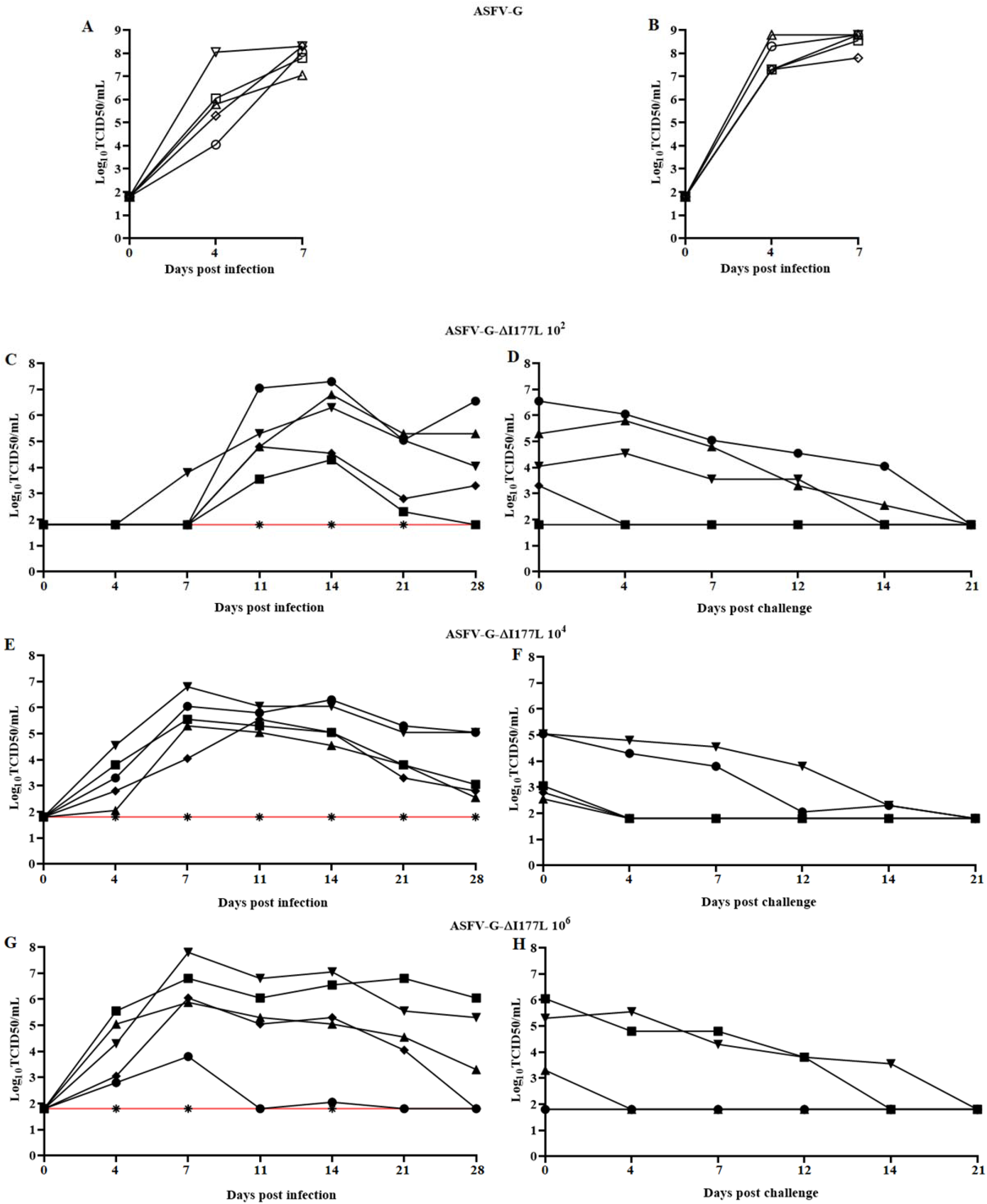
Viremia titers detected in pigs IM inoculated with either **(A)** 10^2^, 10^4^, or 10^6^ HAD_50_ of ASFV-Georgia-ΔI177L or 10^2^ HAD_50_ of ASFV-G. **(B)** Viremia after the challenge with 10^2^ HAD_50_ of ASFV-G Each curve represents values from individual animals in each of the group. Sensitivity of virus detection: ≥ 1.8 log_10_ HAD_50_/ml_50_/ml. Data from sentinel animals are depicted in red.

### Protective efficacy of ASFV-G-ΔI177L against challenge with parental ASFV-G

To assess the ability of ASFV-G-ΔI177L infection to induce protection against challenge with highly virulent parental virus ASFV-G, all animals infected with ASFV-G-ΔI177L were challenged 28 days later with 10^2^ HAD_50_ of ASFV-G by IM route. Five naïve animals were challenged as a mock-inoculated control group.

All mock animals started showing disease-related signs by 3-4 days post challenge (dpc), with rapidly increasing disease severity in the following hours and being euthanized by 5-6 dpc (Table 3). On the other hand, the three groups of animals infected with ASFV-G-ΔI177L remained clinically healthy, not showing any significant signs of disease during the 21-day observational period. Therefore, ASFV-G-ΔI177L-treated animals are protected against clinical disease when challenged with the highly virulent parental virus.

**Table 3.** Swine survival and fever response in ASFV-G-ΔI177L-infected animals challenged with ASFV-G virus 28 days later.

| Virus | No. of survivors/<br>total | Mean time to death<br>(days ± SD) | Fever |  |  |
| --- | --- | --- | --- | --- | --- |
|  |  |  | No. of days to onset<br>(days ± SD) | Duration<br>No. of days<br>(days ± SD) | Maximum daily temp<br>(°F ± SD) |
| Mock | 0/5 | 5.6 (0.55) <sup>(1)</sup> | 4.2 (0.84) | 1.4 (0.88) | 105.6 (0.78) |
| ASFV-G-ΔI177L 10<sup>2</sup> HAD<sub>50</sub> | 10/10 | - | - | - | 102.7 (0.68) |
| ASFV-G-ΔI177L 10<sup>4</sup> HAD<sub>50</sub> | 5/5 | - | - | - | 102.9 (0.37) |
| ASFV-G-ΔI177L 10<sup>6</sup> HAD<sub>50</sub> | 5/5 | - | - | - | 103 (0.43) |

(1) All animals were euthanized due to humanitarian reasons following the corresponding IACUC protocol.

Analysis of viremia in animals infected with ASFV-G presented with expected high titers (10^7.3^ - 10^8.3^ HAD_50_/ml) on day 4 pi, increasing (averaging 10^8.5^ HAD_50_/ml) by the time when all animals were euthanized. After challenge, none of the ASFV-G-ΔI177L-infected animals had viremias with values higher than those present at challenge and viremia values decreased progressively until the end of the experimental period (21 days after challenge) when, importantly, no circulating virus could be detected in blood from any of these animals (Fig. 7). Interestingly, post-challenge viremia titers, calculated by HA, exactly coincide with those calculated by fluorescence suggesting a lack (or at least a very low rate) of replication by the challenge virus. To assess the potential replication of the challenge virus the presence of ASFV-G was tested in blood samples taken at day 4 post challenge, when the highest viremia titers occur after challenge (Fig. 7). Using an I177L-specific real-time PCR to detect only challenge virus (with a demonstrated sensitivity of approximately 10 HAD_50_) all blood samples tested negative but one from an animal infected with 10^2^ HAD_50_ of ASFV-G-ΔI177L (data not shown). Furthermore, tonsils and spleen samples were obtained from all ASFV-G-ΔI177L infected animals at the end of the observational period (21 days post-challenge) and tested for the presence of virus (detected by hemoadsoption) using swine macrophage cultures. Most of the animals in each group had infectious virus either in tonsils or spleen (data not shown). All positive samples were then assessed using the I177L-specific real-time PCR, detecting the presence of the challenge virus in only one spleen belonging to the same animal initially infected with 10^2^ HAD_50_/ml of ASFV-G-ΔI177L, which also had challenge virus in the blood (data not shown). These results suggest that replication of challenge virus was absent in all infected animals receiving 10^4^ HAD_50_/ml or higher and most of the animals receiving 10^2^ HAD_50_/ml of ASFV-G-ΔI177L.

### Host antibody response in animals infected with ASFV-G-ΔI177L

Host immune mechanisms mediating protection against virulent strains of ASFV in animals infected with attenuated strains of virus are not well identified (20–22). Our previous experience indicated that the only parameter consistently associated with protection against challenge is the level of circulating antibodies (18). In order to gain additional understanding of immune mechanisms in ASFV-G-ΔI177L-infected animals, we attempted to correlate the presence of anti-ASFV circulating antibodies with protection. ASFV-specific antibody response was detected in the sera of these animals using two in-house developed direct ELISAs (18). All animals infected with ASFV-G-ΔI177L, regardless of the dose of virus received, possessed similar high titers of circulating anti-ASFV antibodies (Fig. 8). Antibody response, mediated by IgM and IgG isotypes, was detected in all three groups by day 12 pi. By day 14 pi response mediated by both antibody isotypes reached maximum levels in all groups. IgM-mediated antibody response disappeared in all animals by day 21 pi, while IgG mediated response remained high with minimal fluctuation until day 28 pi without significant differences between animals in the three groups inoculated with ASFV-G-ΔI177L. Therefore, as described in our previous reports (14, 18) there is a close correlation between presence of anti-ASFV antibodies at the moment of challenge and protection. It should be mentioned that no antibodies were detected in any serum sample obtained from the sentinel animals corroborating the virological data indicating that sentinel animals were not infected from ASFV-G-ΔI177L infected animals in any of the three groups (data not show).

**Fig. 8:**
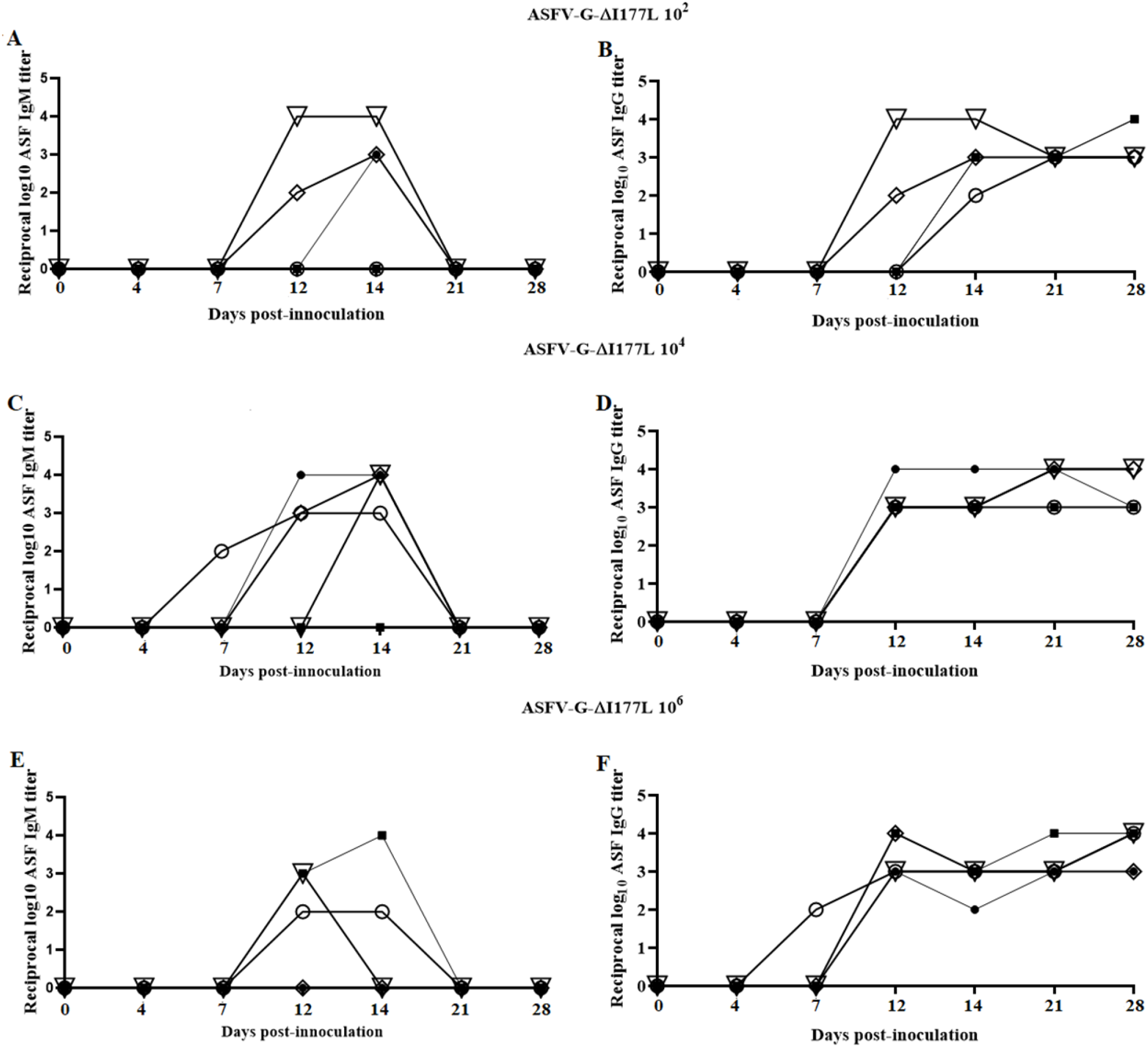
Anti-ASFV antibody (IgM mediated shown in panels **A**, **C** and **E**, and IgG mediated shown in panels **B**, **C** and **F**) titers detected by ELISA in pigs IM inoculated with either 10^2^, 10^4^, or 10^6^ HAD_50_ of ASFV-Georgia-ΔI177L. Each curve represents values from individual animals in each group.

## Discussion

The use of attenuated strains is currently the most plausible approach to develop an effective ASF vaccine. Rational development of attenuated strains by genetic manipulation is a valid alternative, and perhaps safer methodology, compared to the use of naturally attenuated isolates. Several attenuated strains, obtained by genetic manipulation consisting of deletions of single genes or a group of genes, have been shown to induce protection against the virulent parental virus (10–13, 15, 23). Here, the identification of a previously uncharacterized ASFV gene, I177L, as a viral genetic determinant of virulence is described. Deletion of I177L completely attenuates ASFV-G in swine, even when used at doses as high as 10^6^ HAD_50_. Only two other genetic modifications have been shown to completely abolish virulence in the highly virulent ASFV Georgia isolate: deletion of the 9GL gene (particularly potentiated by the additional deletion of the UK gene) and deletion of a group of six genes from the MGF360 and 530 (12, 13, 24). The attenuation observed by deleting the I177L gene is a remarkable discovery since ASFV-G has not been efficiently attenuated by deletion of any other genes that have been associated with attenuation in other ASFV isolates (13, 25). Based on the cumulative efforts supported by several studies, it is apparent that the genetic background where deletion is operated plays a critical role in the effect of a particular gene in virus virulence supporting the concept that AFV virulence is the result of the interactive effect of several virus genes.

Although ASFV-G-ΔI177L-infected animals remained clinically normal, all of them presented with viremia by 28 days pi, in some cases with relatively high titers. Interestingly, no infectious virus or virus specific antibodies could be detected in any of the three sentinels indicating that transmission of ASFV-G-ΔI177L from infected to naïve animals is not a frequent event, a desirable characteristic for a potential candidate live attenuated vaccine.

Importantly, animals infected with ASFV-G-ΔI177L were effectively protected when challenged at 28 dpi. Protection was achieved with doses as low as 10^2^ HAD_50_ of ASFV-G-ΔI177L while even the administration of 10^6^ HAD_50_ ASFV-G-ΔI177L did not produce any disease-associated signs (not even a transient rise in body temperature), emphasizing the safety of ASFV-G-ΔI177L as a potential vaccine candidate. Importantly, it appears that replication of the challenge virus in the ASFV-G-ΔI177L-infected animals is quite restricted since challenge virus was isolated from only one of the animals inoculated with the low dose of ASFV-G-ΔI177L.

Although the host mechanisms mediating protection against ASFV infection remain under discussion (1, 2), in our experience with different live attenuated vaccine candidates we have been observing a close association between presence of circulating virus-specific antibodies and protection (12–14, 24, 26). In this report, we were also able to associate presence of virus specific antibodies and protection. Interestingly, regardless of the ASFV-G-ΔI177L dose used, all animals had similar antibody titers at the time of challenge, supporting the fact that low doses of ASFV-G-ΔI177L were as effective as the highest dose. A note is the fact that by day 14 pi, all animals reached maximum antibody titers. Although in this report challenge was not performed at 14 dpi this data agrees with previous published reports demonstrating that animals inoculated with vaccine candidate ASFV-G-Δ9GL/ΔUK presenting with circulating antibodies were protected against challenge at 2 weeks post-infection ^24^.

We believe results presented here demonstrate that ASFV-G-ΔI177L can be considered a strong vaccine candidate to protect animals against the ASFV Georgia isolate and its derivatives currently causing outbreaks in a wide geographical area from central Europe to China and Southeast Asia. The complete lack of residual virulence, even when administered at high doses, apparent low levels of transmissibility to naïve animals, and its high efficacy in inducing protection even at low doses makes ASFV-G-ΔI177L a promising novel vaccine candidate.

## Acknowledgements

We thank the Plum Island Animal Disease Center Animal Care Unit staff for excellent technical assistance. We wish to, particularly, thank Mrs. Melanie V. Prarat for editing the manuscript. This research was supported in part by an appointment to the Plum Island Animal Disease Center (PIADC) Research Participation Program administered by the Oak Ridge Institute for Science and Education (ORISE) through an interagency agreement between the U.S. Department of Energy (DOE) and the U.S. Department of Agriculture (USDA). ORISE is managed by ORAU under DOE contract number DE-SC0014664. All opinions expressed in this paper are the author’s and do not necessarily reflect the policies and views of USDA, ARS, APHIS, DHS, DOE, or ORAU/ORISE.

## References

1. Tulman ER, Delhon, G.A., Ku, B.K. and Rock, D.L.. 2009. African Swine Fever Virus p43–87, Lesser Known Large dsDNA Viruses, vol 328 Springer-Verlag Berlin Heidelberg.

2. Costard S, Porphyre, V., Messad, S., Rakotondrahanta, S., Vidon, H., Roger, F. and Pfeiffer, D. U. Exploratory multivariate analysis for differentiating husbandry practices relevant to disease risk for pig farmers in Madagascar, p 228–238. *In* (ed),

3. Chapman DA, Darby AC, Da Silva M, Upton C, Radford AD, Dixon LK. 2011. Genomic analysis of highly virulent Georgia 2007/1 isolate of African swine fever virus. Emerg Infect Dis 17:599–605.

4. Coggins L. 1974. African swine fever virus. Pathogenesis. Prog Med Virol 18:48–63.

5. Forman AJ, Wardley RC, Wilkinson PJ. 1982. The immunological response of pigs and guinea pigs to antigens of African swine fever virus. Arch Virol 74:91–100.

6. Kihm UAM, Mueller H, Pool R. 1987. Approaches to vaccination African swine Fever. Martinus Nijhoff Publishing, Boston.

7. Mebus CA. 1988. African swine fever. Adv Virus Res 35:251–69.

8. Hamdy FM, Dardiri AH. 1984. Clinical and immunologic responses of pigs to African swine fever virus isolated from the Western Hemisphere. Am J Vet Res 45:711–4.

9. Ruiz-Gonzalvo F CM, Bruyel V. 1981. Immunological responses of pigs to partially attenuated ASF and their resistance to virulent homologous and heterologous viruses., Rome.

10. Lewis T, Zsak L, Burrage TG, Lu Z, Kutish GF, Neilan JG, Rock DL. 2000. An African swine fever virus ERV1-ALR homologue, 9GL, affects virion maturation and viral growth in macrophages and viral virulence in swine. J Virol 74:1275–85.

11. Moore DM, Zsak L, Neilan JG, Lu Z, Rock DL. 1998. The African swine fever virus thymidine kinase gene is required for efficient replication in swine macrophages and for virulence in swine. J Virol 72:10310–5.

12. O’Donnell V, Holinka LG, Gladue DP, Sanford B, Krug PW, Lu X, Arzt J, Reese B, Carrillo C, Risatti GR, Borca MV. 2015. African Swine Fever Virus Georgia Isolate Harboring Deletions of MGF360 and MGF505 Genes Is Attenuated in Swine and Confers Protection against Challenge with Virulent Parental Virus. J Virol 89:6048–56.

13. O’Donnell V, Holinka LG, Krug PW, Gladue DP, Carlson J, Sanford B, Alfano M, Kramer E, Lu Z, Arzt J, Reese B, Carrillo C, Risatti GR, Borca MV. 2015. African Swine Fever Virus Georgia 2007 with a Deletion of Virulence-Associated Gene 9GL (B119L), when Administered at Low Doses, Leads to Virus Attenuation in Swine and Induces an Effective Protection against Homologous Challenge. J Virol 89:8556–66.

14. O’Donnell V, Holinka LG, Sanford B, Krug PW, Carlson J, Pacheco JM, Reese B, Risatti GR, Gladue DP, Borca MV. 2016. African swine fever virus Georgia isolate harboring deletions of 9GL and MGF360/505 genes is highly attenuated in swine but does not confer protection against parental virus challenge. Virus Res 221:8–14.

15. Zsak L, Lu Z, Kutish GF, Neilan JG, Rock DL. 1996. An African swine fever virus virulence-associated gene NL-S with similarity to the herpes simplex virus ICP34.5 gene. J Virol 70:8865–71.

16. Reed LJM, H. 1938. A simple method of estimating fifty percent endpoints. The American Journal of Hygiene 27:493–497.

17. Borca MV, Holinka LG, Berggren KA, Gladue DP. 2018. CRISPR-Cas9, a tool to efficiently increase the development of recombinant African swine fever viruses. Sci Rep 8:3154.

18. Carlson J, O’Donnell V, Alfano M, Velazquez Salinas L, Holinka LG, Krug PW, Gladue DP, Higgs S, Borca MV. 2016. Association of the Host Immune Response with Protection Using a Live Attenuated African Swine Fever Virus Model. Viruses 8.

19. Forth JH, Forth LF, King J, Groza O, Hubner A, Olesen AS, Hoper D, Dixon LK, Netherton CL, Rasmussen TB, Blome S, Pohlmann A, Beer M. 2019. A Deep-Sequencing Workflow for the Fast and Efficient Generation of High-Quality African Swine Fever Virus Whole-Genome Sequences. Viruses 11.

20. Onisk DV, Borca MV, Kutish G, Kramer E, Irusta P, Rock DL. 1994. Passively transferred African swine fever virus antibodies protect swine against lethal infection. Virology 198:350–4.

21. Ruiz Gonzalvo F, Carnero ME, Caballero C, Martinez J. 1986. Inhibition of African swine fever infection in the presence of immune sera in vivo and in vitro. Am J Vet Res 47:1249–52.

22. Oura CA, Denyer MS, Takamatsu H, Parkhouse RM. 2005. In vivo depletion of CD8+ T lymphocytes abrogates protective immunity to African swine fever virus. J Gen Virol 86: 2445–50.

23. Zsak L, Caler E, Lu Z, Kutish GF, Neilan JG, Rock DL. 1998. A nonessential African swine fever virus gene UK is a significant virulence determinant in domestic swine. J Virol 72: 1028–35.

24. O’Donnell V, Risatti GR, Holinka LG, Krug PW, Carlson J, Velazquez-Salinas L, Azzinaro PA, Gladue DP, Borca MV. 2017. Simultaneous Deletion of the 9GL and UK Genes from the African Swine Fever Virus Georgia 2007 Isolate Offers Increased Safety and Protection against Homologous Challenge. J Virol 91.

25. Ramirez-Medina E, Vuono E, O’Donnell V, Holinka LG, Silva E, Rai A, Pruitt S, Carrillo C, Gladue DP, Borca MV. 2019. Differential Effect of the Deletion of African Swine Fever Virus Virulence-Associated Genes in the Induction of Attenuation of the Highly Virulent Georgia Strain. Viruses 11.

26. Sanford B, Holinka LG, O’Donnell V, Krug PW, Carlson J, Alfano M, Carrillo C, Wu P, Lowe A, Risatti GR, Gladue DP, Borca MV. 2016. Deletion of the thymidine kinase gene induces complete attenuation of the Georgia isolate of African swine fever virus. Virus Res 213:165–171.

